# Phytohormone production by the arbuscular mycorrhizal fungus *Rhizophagus irregularis*

**DOI:** 10.1101/2020.06.11.146126

**Authors:** Simon Pons, Sylvie Fournier, Christian Chervin, Guillaume Bécard, Soizic Rochange, Nicolas Frei Dit Frey, Virginie Puech Pagès

## Abstract

Arbuscular mycorrhizal symbiosis is a mutualistic interaction between most land plants and fungi of the glomeromycotina subphylum. The initiation, development and regulation of this symbiosis involve numerous signalling events between and within the symbiotic partners. Among other signals, phytohormones are known to play important roles at various stages of the interaction. During presymbiotic steps, plant roots exude strigolactones which stimulate the fungus, and favour the initiation of symbiosis. At later stages, different plant hormone classes can act as positive or negative regulators of the interaction. Although the fungus is known to reciprocally emit regulatory signals, its potential contribution to the phytohormonal pool has received little attention, and has so far only been addressed by indirect assays. In this study, using mass spectrometry, we analyzed phytohormones released into the medium by germinated spores of the arbuscular mycorrhizal fungus *Rhizophagus irregularis*. We detected the presence of a cytokinin (isopentenyl-adenosine) and an auxin (indole-acetic acid). In addition, we identified a gibberellin (gibberellic acid 4) in spore extracts. We also used gas chromatography to show that *R. irregularis* produces ethylene from methionine and the α-keto γ-methylthiobutyric acid pathway. These results highlight the possibility for AM fungi to use phytohormones to interact with their host plants, or to regulate their own development.

## Introduction

Arbuscular Mycorrhizal (AM) symbiosis is a 460 million-year-old interaction [1] between glomeromycotina fungi and over 70% of land plants [2]. In angiosperms, AM fungi colonize the inner root cortex of their host to develop intracellular ramified structures called arbuscules. These arbuscules are the main site for nutrient exchange between the plant and the fungus. AM fungi provide their host plant with water and minerals, and in return receive carbon sources (mainly sugars and lipids) [3,4]. As AM fungi are obligate biotrophs, this interaction is essential for their growth, development and reproduction. On the plant side, this interaction improves nutrition and resistance to biotic and abiotic stresses [5–7].

Prior to physical contact, the two partners of the AM symbiosis interact via signalling molecules [8,9]. Host roots release several types of compounds affecting the presymbiotic development of AM fungi, such as some flavonoids, phenolic compounds, hydroxy fatty acids [10,11]. Particular attention has been paid to the root-exuded strigolactone phytohormones: they stimulate the germination of AM fungal spores, the oxidative metabolism and branching of germinating hyphae, and finally root colonization [12–15]. In addition, *N*-acetylglucosamine-based compounds could be exchanged in both directions: lipochitooligosaccharides (LCOs) and chitooligosaccharides (COs) are released by germinating spores of AM fungi and stimulate the initiation of the symbiosis [16–18], and a plant exporter of N-acetylglucosamine has been shown to be required for the first steps of the interaction [19]. Finally, additional, yet unidentified, signals of plant or fungal origin may act prior to root colonization [20–23].

Later stages of AM interactions are regulated by a number of factors, including nutrient exchange [23] and phytohormones [24,25]. Analysis of AM symbiosis regulation by phytohormones has revealed a complex pattern of modified hormonal contents or altered response to hormones in mycorrhizal plants, and reciprocal effects of exogenous hormone application on the symbiotic interaction. Although observations were made across a wide range of combinations of plant/fungal species and experimental conditions, it is possible to draw broad conclusions about the role of the different hormone families. Auxins (AUX), abscisic acid (ABA) and brassinosteroids (BRs) have been identified as positive regulators of the AM symbiosis [26–28]. On the contrary, gibberellins (GAs) and salicylic acid (SA) have been described as negative regulators of the interaction [29–31]. The effects of cytokinins have not yet been clearly established [32]. Finally, the role of ethylene (ET) and jasmonic acid (JA) seems to vary with their concentration [33–36].

Importantly, these studies have addressed the role of phytohormones in the AM symbiosis by two main approaches: the analysis of plant mutants affected in phytohormone synthesis or perception, or the treatment of mycorrhizal plants with exogenous hormones. The study of hormone perception mutants clearly addresses the effects of hormones on the plant. In contrast, both exogenous treatments and hormone deficiency in the plant result in modified hormonal contents in colonized roots, which could impact either or both symbionts. In spite of this, and because phytohormones are generally seen as plant signals, results of such studies are commonly interpreted exclusively in terms of impacts on the plant. Likewise, changes in hormonal contents measured in mycorrhizal plants are usually attributed to modifications of hormonal metabolism in plant cells. This interpretation ignores a potential contribution of the fungal partner to the hormonal pool. Yet, many microorganisms can produce phytohormones and this could also be the case of AM fungi. Among soil microorganisms interacting with plants, plant growth-promoting rhizobacteria and fungi have been shown to produce auxin, cytokinins, ABA and gibberellins [37–39], and this can contribute to their growth-promoting effects [40]. Nitrogen-fixing rhizobia associated with legumes also produce a range of phythormones [41]. In the fungal kingdom, phytohormone production has been documented in symbionts like ectomycorrhizal fungi [42,43], as well as in pathogens [44]. Ethylene is quite common among phytohormones produced by fungi [45–48] and in some cases the biosynthetic pathways have been characterized. The α-keto γ-methylthiobutyric acid (KMBA) pathway, well described in *Botrytis cinerea* [47], requires the deamination of methionine into KMBA. Subsequently, KMBA can be oxidised into ethylene through different means. It can either be oxidized by hydroxyl radicals [49], peroxidases [50], or in a light-dependant manner, induced by the photo-oxidation of flavins [51], leading to the production of ethylene. In contrast, the second ethylene-producing pathways in microorganisms involves a very specific enzyme. The Ethylene Forming Enzyme (EFE), described in *Penicillium digitatum*, or *Fusarium oxysporum* [45,52], produces ethylene through two simultaneous reactions using L-arginine and 2-oxoglutarate as co-substrates. Both pathways differ from the main pathway for ethylene production in plants, which involves a light-independent and methionine-dependent pathway requiring the aminocyclopropane-carboxylate (ACC) synthase (ACS) and ACC oxidase (ACO).

The fact that many plant-associated microorganisms produce phytohormones raises the possibility of a similar behaviour in AM fungi which have co-evolved with plants for over 400 million years. This question is challenging to address experimentally, essentially because of the obligate biotrophy of these fungi. This feature implies that isolated fungi can only be kept in culture for short periods of time, and limits the availability of biological material. Nevertheless, previous studies have provided indirect evidence for the presence of phytohormones in AM fungi. ELISA tests indicated that spores and hyphae of *Rhizophagus* (formerly *Glomus*) species could contain aglycone and glycosylated ABA [53], while indirect bioassays suggested the presence of gibberellin-like and cytokinin-like molecules [54]. A direct analysis by GC-MS of spore extracts revealed the presence of small amounts of IAA in *Glomus intraradices*, but in this study spores were directly taken from maize pot cultures, hence not in axenic conditions, and may have been contaminated with root fragments or other microorganisms [55]. Genes encoding a CLAVATA3/Embryo surrounding region-related (CLE) peptide hormone proposed to positively regulate the symbiosis have also been identified in AM fungal genomes [56].

In this study, we analysed the presence of phytohormones in germinating spores, or in their exudates, of the model AM fungus *Rhizophagus irregularis* grown axenically. We used a combination of gas/liquid chromatography and mass spectrometry to allow unambiguous compound identification. In the case of ethylene, we investigated the putative biosynthetic pathways through the use of metabolic precursors.

## Materials & methods

### Chemicals, reagents and standards

Phytohormone standards were purchased from Olchemim (iP, iPR, iP9G, Ki, *m*T, *t*Z, *c*Z, *t*ZR, *c*ZR, DHZ, BAP, GA_1_, IAA-Asp, JA-Ile, ABA-GE, BL), Fluka (IBA, NAA), Acros Organics (IAA, IPA, SA, ABA), Sigma-Aldrich (IAA-Ala, JA, MeJA, Strigol), Fisher chemical (GA_3_) and Duchefa (GA_4_). We prepared the standards following the manufacturer’s recommendations, and stored solutions at −20 °C. L-Methionine and α-keto-γ-methylthiobutyrate (KMBA) were purchased from Sigma-Aldrich. LC/MS-grade acetonitrile and HPLC-grade methanol were purchased from Fisher Chemical, formic acid from Acros Organics, and ammonium hydroxide from Sigma-Aldrich.

### Fungal culture and exudate preparation

*Rhizophagus irregularis* DAOM 197198 sterile spores were purchased from Agronutrition (Labège, France). The spores were produced in axenic conditions. Spore numbers were determined by the supplier by counting spores in an aliquot of the sold suspension with a binocular microscope. Spores were rinsed from their storage buffer using a 40 µm nylon cell strainer (VWR) by five washes with sterile UHQ water. Spores were resuspended in sterile UHQ water and stored at 4 °C before use.

For the production of germinated spore exudates (GSE), spores were germinated in sterile UHQ water in a CO_2_ incubator (30 °C, 2 % CO_2_) for 7 days with a concentration of 400 spores.mL^−1^ in 25 mL Petri dishes. GSE were filtered through a glass-fiber frit (Chromabond, Macherey-Nagel, France), then frozen in liquid nitrogen and freeze-dried. Filtered spores were collected and stored at −80 °C.

### Phytohormone and KMBA extraction

#### From germinated spore exudates

freeze-dried GSE from 10,000 spores or 250,000 spores of *R. irregularis* were reconstituted in 100 µL of 1 M formic acid and stored at −20°C before MS analysis.

#### From ground spores

the protocol of phytohormone extraction and separation by Solid Phase Extraction (SPE) was adapted from Kojima *et al.* [57] as follows. Two hundred and fifty thousand spores were hand-ground in liquid nitrogen with a mortar and pestle, resuspended in 1 mL of cold modified Bieleski’s solvent (methanol/water/formic acid 75:20:5, v:v:v; [58]) and left overnight at −20 °C to achieve complete extraction. The crude extracts were centrifuged for 15 min at 10,000 x *g*, at 4 °C. The pellet was reextracted in 200 µL modified Bieleski’s solvent for 30 min at −20 °C, centrifuged 15 min at 10,000 x *g*, and the supernatant pooled with the first one. Extracts were pre-purified on SPE Oasis HBL cartridges (1 mL per 30 mg, Waters). Cartridges were conditioned in 1 mL methanol and equilibrated in 1 mL of 1 M formic acid. Two-mL samples were loaded and eluted with 1 mL modified Bieleski’s solvent. Eluates containing phytohormones were dried under nitrogen stream and reconstituted in 1 mL of 1 M formic acid. The separation of phytohormones was achieved with SPE Oasis MCX cartridges (1 mL per 30 mg, Waters). Cartridges were conditioned in 1 mL methanol and equilibrated in 1 mL of 1 M formic acid. Samples were successively eluted by 1 mL methanol (fraction 1, containing neutral and acidic hormones), 1 mL of 0.35 M ammonium hydroxide (fraction 2) and 1 mL methanol/0.35 M ammonium hydroxide 60:40 (fraction 3, containing cytokinins) [57]. The three fractions were dried under nitrogen stream and kept at 4 °C before analysis. Fractions were reconstituted in 100 μL 1 M formic acid and then analysed by LC-MS.

### LC-MS analysis of phytohormones

Samples (GSEs or fractions obtained from ground spores as previously described) were analysed by UHPLC-MS with two types of mass spectrometers, a Q-Trap 5500 (AB Sciex) in MRM mode for higher detection sensitivity, and a Q-Exactive Plus™ (Thermo Scientific) for higher mass accuracy by high resolution analysis.

Twenty-one biological replicates of GSE and 12 replicates of ground spores were used to perform phytohormone detection. IPR was consistently identified in GSE, while IAA was detected in 76% of them, and thirteen samples were used to perform quantification. GA_4_ was detected in 66% of the ground spore samples.

#### Multiple reaction monitoring (MRM) analysis

A UHPLC system (Dionex Ultimate 3000, Thermo Scientific) was equipped with a Kinetex C18 column (100 × 2.1 mm, 2.6 μm, 100 Å, Phenomenex) heated at 45 °C. Five-μL samples were injected. Separation was performed at a constant flow rate of 0.3 mL.min^−1^, in a gradient of solvent A (water + 0.1% formic acid) and B (acetonitrile + 0.1% formic acid): 1 min 5% B; 11 min 5% to 100% B; 2 min 100% B, and re-equilibration to the initial conditions in 4 min. The Q-Trap 5500 mass spectrometer was equipped with an electrospray source. Curtain gas was set to 30 psi, nebulizer to 40 psi and turbo gas to 60 psi. Capillary voltage was set to 5.5 kV (positive mode) or −4.5 kV (negative mode) on Electrospray Ionization (ESI) source (400 °C). Samples were monitored in positive and negative modes in scheduled Multiple Reaction Monitoring (MRM) mode (60s). Using 25 standards of free or conjugated hormones, in infusion mode (7 µL.min^−1^), the best parameters for declustering potential, collision energy and collision cell exit potential were selected for precursor and product ions measurement. Ionization mode, selected MRM transitions, limit of detection (LOD) and retention time for each hormone are listed in S1 Table. Limits of detection and quantification were determined using standards diluted from 0.1 mM to 1pM in methanol. By this approach, we could perform an approximate quantification of phytohormones in fungal samples. Data processing was performed using Analyst 1.6.2 software.

#### High resolution mass spectrometry (HRMS) analysis

A UHPLC system (Ultimate 3000 RSLC system, Thermo Scientific) was equipped with a Hypersil Gold aQ C18 column (100 mm x 2.1 mm, 1.9 μm, 175 Å, Thermo Scientific #25302102130), heated at 35°C. Five-μL samples were injected. Separation was performed at a constant flow rate of 0.3 mL.min^−1^, in a gradient of solvent A (water + 0.05 % formic acid) and B (acetonitrile + 0.05 % formic acid): 1 min 5% B; 7 min 5% to 96% B; 1 min 96% B and re-equilibration to the initial conditions in 3 min. The Q-Exactive Plus™ mass spectrometer was equipped with a H-ESI II probe, heated at 256 °C. Sheath gas was set to 48, sweep gas to 2, auxiliary gas to 11, and heated at 413 °C. Capillary voltage was set to 3.5 kV in positive mode and −2.5 kV in negative mode. Ionization was performed in positive and negative modes, in full scan analysis (centroid), with a resolution of 35,000. Automatic Gain Control was set to 3.10^6^, with a 50 to 600 *m/z* scan range. A Target-MS/MS scan of confirmation of the phytohormone, based on the specified inclusion list (5 ppm), was triggered when the mass spectrometer detected a known phytohormone in an MS spectrum. In this case, Automatic Gain Control was set to 2.10^5^, and resolution to 17,500. Data processing was performed by Xcalibur 3.0 and Tracefinder 3.2 softwares.

### Ethylene detection

20,000 spores in 2 mL of sterile UHQ water, either untreated, or supplemented with 10 mM of methionine or 1 µM of KMBA, were incubated in a sterilized glass tube (Ø=1.35 cm, H= 6.10 cm, V=7.5 mL, dead volume = 5.5 mL) wrapped in tinfoil to avoid light exposure, and sealed with a porous silicone stopper (Hirschmann Instruments). They were germinated for three days in a CO_2_ incubator (30°C, 2% CO_2_). The stopper was then replaced by an air-tight rubber stopper, and spores were confined for 1 day and exposed or not to light (100 μmol photons m^−2^ s^−1^, 21°C).

The headspace ethylene content was assayed by gas chromatography as described previously [59]. One mL of headspace gas was manually injected into a GC-FID (Agilent 7820a), equipped with a 80/100 alumina column (1/8” x 2 mm x 1.5 m, Agilent) and set with the following parameters: oven temperature 70 °C, injector temperature 110 °C, N_2_ vector gas flow rate 28 mL.min^−1^, flame ionization detector temperature 250 °C. Ethylene peak area was measured and normalized with the O_2_ injection peak area. Its retention time and calibration were determined with an external standard of 4 ppm of ethylene.

For ethylene production assays: darkness without spores n=13, light without spores n=16, darkness with spores n=17, light with spores n=18, light and methionine without spores n=7, darkness and methionine with spores n=9, light and methionine with spores n=24, light and KMBA without spores n=6, light and KMBA with spores n=6.

### KMBA detection by LC-MS

A UHPLC system (Dionex Ultimate 3000, Thermo Scientific) was equipped with a reverse-phase column Acquity UPLC BEH-C18 (2.1 × 150 mm, 1.7 μm, Waters) heated at 45 °C. Ten-μL samples were injected. Separation was performed at a constant flow rate of 0.3 mL.min^−1^, in a gradient of solvent A (water + 0.1% formic acid) and B (acetonitrile): 1 min 5% B; 8 min 5% to 100% B; 2 min 100% B, and re-equilibration to the initial conditions in 2 min.

A Q-Trap 4500 mass spectrometer (AB Sciex) was used with an electro-spray ionization source in the negative ion mode. Curtain gas was set to 30 psi, nebulizer to 40 psi and turbo gas to 60 psi. Capillary voltage was set to −3.5 kV (negative mode) on Electrospray Ionization (ESI) source (400 °C). Optimizations of the source parameters were done using the KMBA standard at 10^−5^ M water by infusion at 7 μL.min^−1^, using a syringe pump. Three GSE sample were analysed for each condition. Data processing was performed using Analyst 1.6.2 software.

### Sequence analysis

Glomeromycotina (taxid:214504) nucleotide and protein sequences were analysed using TBLASTN and BLASTP searches with default parameters on the NCBI website. The 2-oxoglutarate-dependent ethylene/succinate-forming enzyme from *Penicillium digitatum* XP_014538251.1 was chosen as query to identify putative EFEs in *R. irregularis*.

### Statistical analysis

The version 4.0.0 of R, with the version 1.3-1 of package Agricolae and the version 3.1.0 of package GGPlot2 were used for statistical analysis. In order to compare ethylene production between all groups, a non-parametric analysis was carried out using Kruskal-Wallis test and pairwise comparisons were performed using FDR adjustment for multiple comparisons.

## Results

### Detection of phytohormones in germinated spore exudates and germinated spore extracts

The aim of this study was to investigate the production of phytohormones by the model AM fungus *R. irregularis*. To avoid any contamination with plant-borne phytohormones, it was crucial to start from pure fungal material. We used spores of *R. irregularis* produced in axenic conditions, and the spores, free of root debris, were carefully rinsed to eliminate the storage solution.

We started with an analysis of exudates produced by *R. irregularis* spores germinated in water for seven days. These Germinated Spore Exudates (GSE) were concentrated and analysed by Liquid Chromatography (LC) coupled to Mass Spectrometry (MS). The detection of a total of 38 compounds, covering eight hormone families (S1 Table), was attempted. Synthetic standards were available for 25 of these compounds, allowing direct comparison of retention times and MS data. Two types of MS analyses were successively carried out. The highly sensitive Multiple Reaction Monitoring (MRM) mode was used to look for characteristic precursor-to-product-ion *m/z* transitions upon fragmentation. The retention times associated with these *m/z* transition signals were compared with the retention times of corresponding standards. To further ascertain hormone identification, High-Resolution Mass Spectrometry (HRMS) was then used to extract signals for precursor and product ions of the expected accurate *m/z* (+/− 5 ppm), and again was performed in comparison with standards.

Using these two approaches, we identified two hormones in GSE samples produced by 10,000 spores: the cytokinin isopentenyl-adenosine (iPR) and the auxin indole-acetic acid (IAA) (Fig 1A and D). In the MRM mode, iPR was detected with the *m/z* transitions 336 > 204 and 336 > 136; IAA was identified with the *m/z* transitions 176 > 130 and 176 > 77 (Fig 1B and E). For both compounds, the observed retention times matched those of the corresponding standards (Fig 1B and E). We were able to detect accurately iPR and IAA in 16 out of 21 biological replicates. The other phytohormones presented in S1 Table were not detected in the GSE samples. According to external standard curves, we estimated that one spore could on average exude 1.2 attomole of iPR (+/− 1.3 attomole) and 29 attomole of IAA (+/− 25 attomole), during seven days of germination. Compound identification was confirmed through HRMS using GSE from 250,000 spores. The precursor ion of *m/z* 336.1662 for iPR, detected at a retention time of 5.78 min (Fig 1C), yielded after selection and fragmentation a product ion of *m/z* 204.1246 (S1 Fig). The precursor ion of *m/z* 176.0706 for IAA, detected at a retention time of 6.18 min (Fig 1F), yielded after selection and fragmentation a product ion of *m/z* 130.0650 (S2 Fig). The mass data, as well as retention times, matched those of the corresponding standards (Fig 1C and F).

**Fig. 1.**
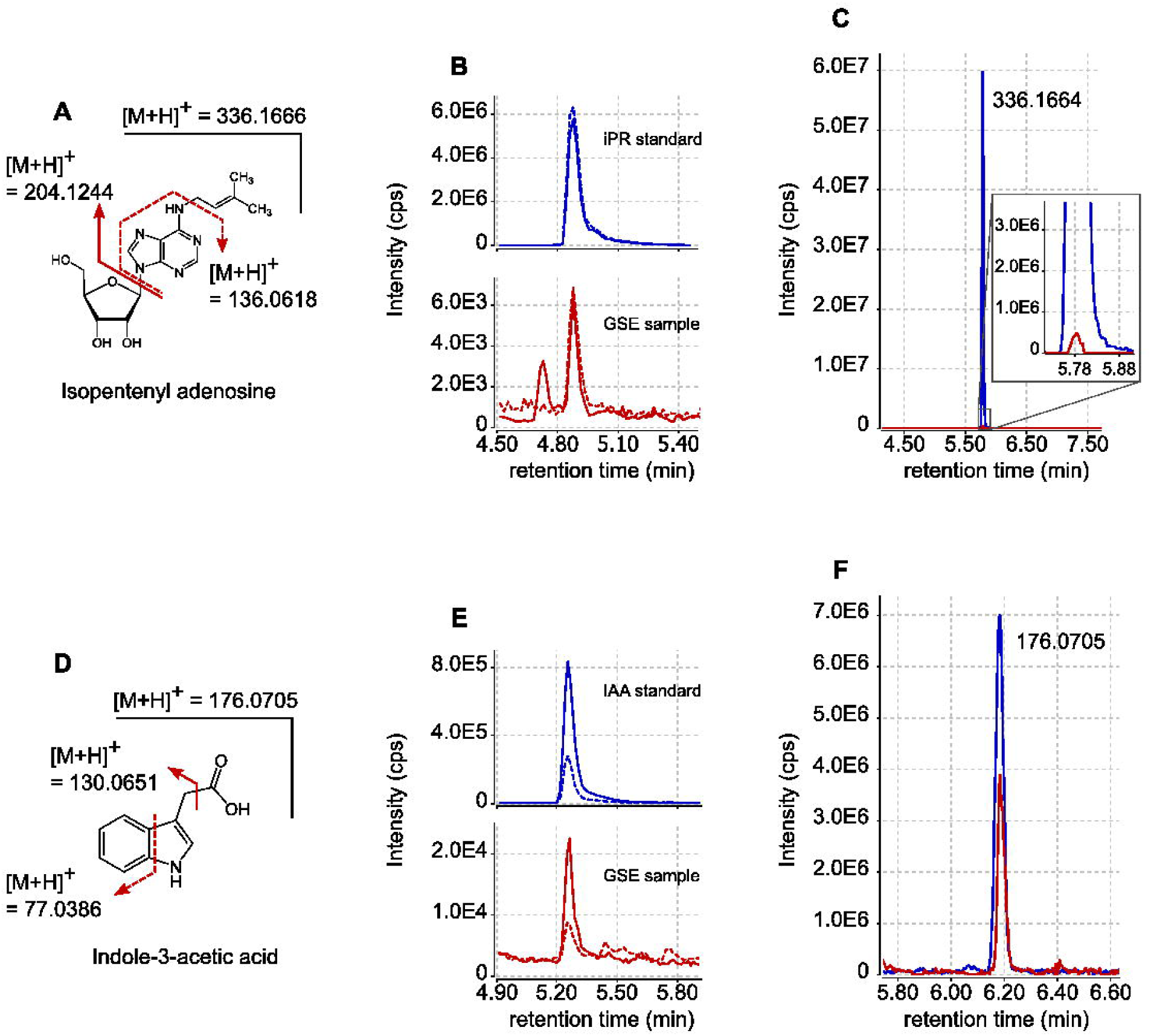
Detection by LC-MS of iPR and IAA exuded by *R. irregularis spores*. (A) Structure and fragmentation pattern of iPR. (B) UPLC-MRM-MS chromatograms of iPR in positive mode. Blue lines represent the signals obtained for iPR external standard (100 nM). Red lines represent the signals obtained with GSE produced by 10,000 *R. irregularis* spores. Plain lines are for *m/z* transition 336 > 204. Dashed lines are for *m/z* transition 336 > 136. (C) LC-HRMS extracted ion chromatogram (XIC) for *m/z* = 336.1666 (+/−5ppm). The blue line represents the signal obtained for iPR external standard (300 nM). The red line represents the signal obtained with GSE produced by 250,000 *R. irregularis* spores. (D) Structure and fragmentation pattern of IAA. (E) UPLC-MRM-MS chromatograms of IAA in positive mode. Blue lines represent the signals obtained for IAA external standard (100 nM). Red lines represent the signals obtained with GSE produced by 10,000 *R. irregularis* spores. Plain lines are for *m/z* transition 176 > 131. Dashed lines are for *m/z* transition 176 > 77. (F) LC-HRMS XIC for *m/ z*= 176.0705 (+/−5ppm). The blue line represents the signal obtained for IAA external standard (300 nM). The red line represents the signal obtained with GSE produced by 250,000 *R. irregularis* spores. Signal intensities are displayed in counts per second (cps).

To investigate whether additional hormones could be present in *R. irregularis* spores, but not released (or in very low amounts) into GSE, we next analysed extracts of 250,000 ground spores. Extracts were pre-purified through two solid-phase extraction (SPE) steps and the fractions where hormones were expected were analysed by LC-MS/MS. This approach allowed the detection in MRM mode of MS-MS transition signals characteristic of a third phytohormone, gibberellic acid 4 (GA_4_). Transitions *m/z* 331 > 257 and 331 > 213 (Fig 2A) were detected in our samples, at almost the same retention time as the standard (retention time shift = −0.09 min, Fig 2B). To investigate whether this slight shift was due to matrix interactions [60], we spiked our sample with the GA_4_ standard. This addition yielded a single chromatographic peak without any splitting, at the same retention time as the spore sample alone and with a doubled intensity. We can therefore attribute the slight retention time difference in Fig 2B to matrix interactions during chromatographic separation.

**Fig. 2.**
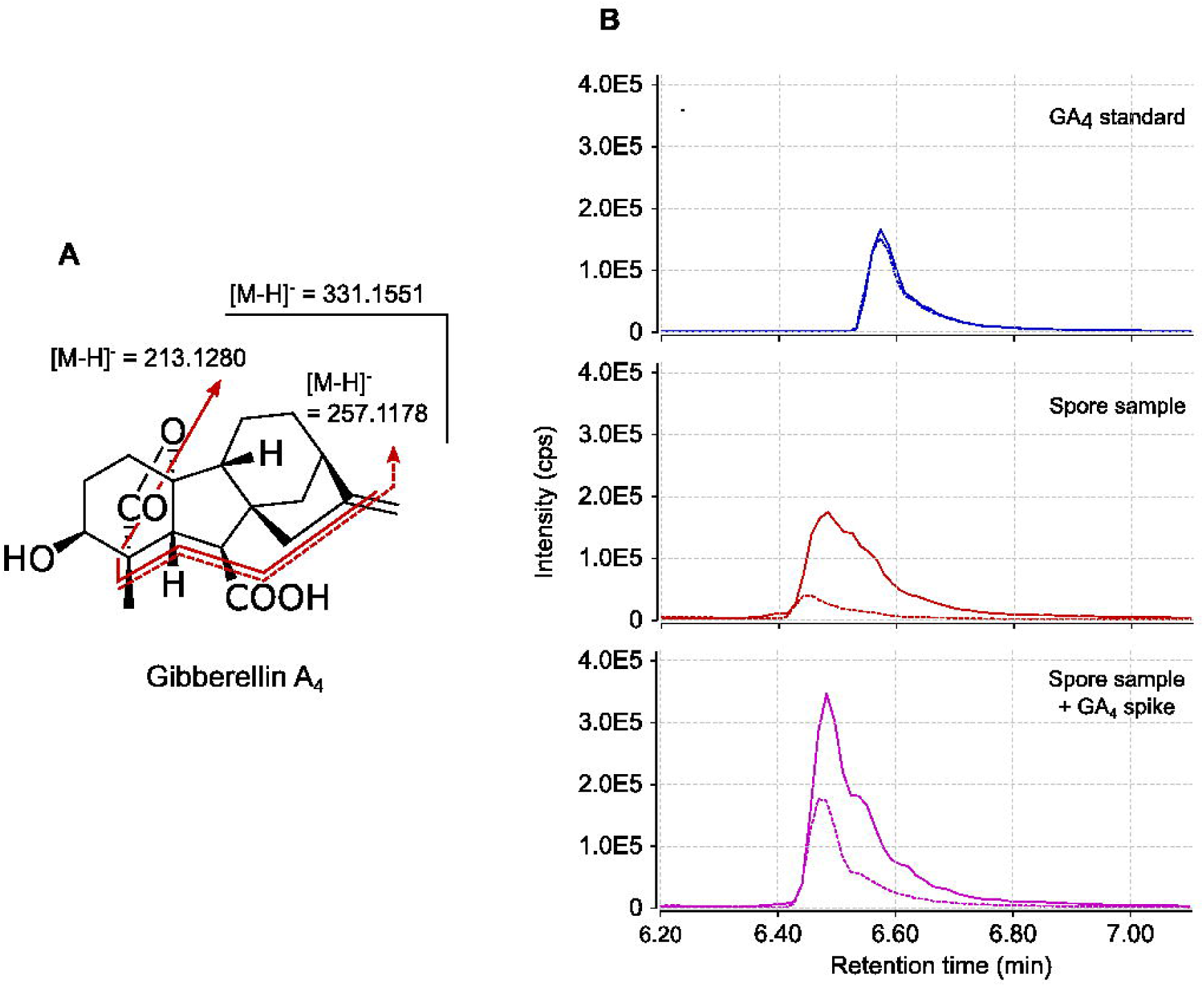
GA_4_ detection in multiple reactions monitoring (MRM) mode. (A) Structure and fragmentation pattern of GA_4_. (B) UPLC-MRM-MS chromatogram of GA_4_ in negative mode. Top (blue lines): External standard (30 nM) of GA_4_. Middle (red lines): pre-purified SPE fraction from 250,000 ground spores of *R. irregularis*. Bottom (purple lines): pre-purified SPE fraction from 250,000 ground spores of *R. irregularis* spiked with GA_4_ standard to a final concentration of 30 nM. Plain lines are for *m/z* transition 331 > 213. Dashed lines are for *m/z* transition 331 > 257. Signal intensities are displayed in counts per second (cps).

### Production of ethylene by *R. irregularis* germinated spores

The release of ethylene by germinating spores was analysed by gas chromatography. To this end, spores were first allowed to germinate for three days in water, in test tubes closed with a gas-permeable stopper. The stopper was then replaced by a gas-tight rubber stopper, and spores were incubated for an additional 24 hours. Gas in the headspace was then sampled for ethylene analysis. As detailed below, light dependency can be used as a criterion to distinguish between ethylene biosynthesis pathways. Therefore, ethylene production was assessed comparatively in spores protected or not from light.

We noticed in control tubes containing no spores that a background quantity of ethylene was produced in the dark as well as in the light (Fig 3, conditions 1 and 2) [61]. The presence of spores slightly enhanced ethylene production in the dark (Fig 3, conditions 1 and 5). When exposed to light, germinating spores produced about 3 times more ethylene than in the dark, suggesting that this production was dependent on the KMBA pathway (Fig 3, conditions 5 and 7) [47,49,51].

**Fig. 3.**
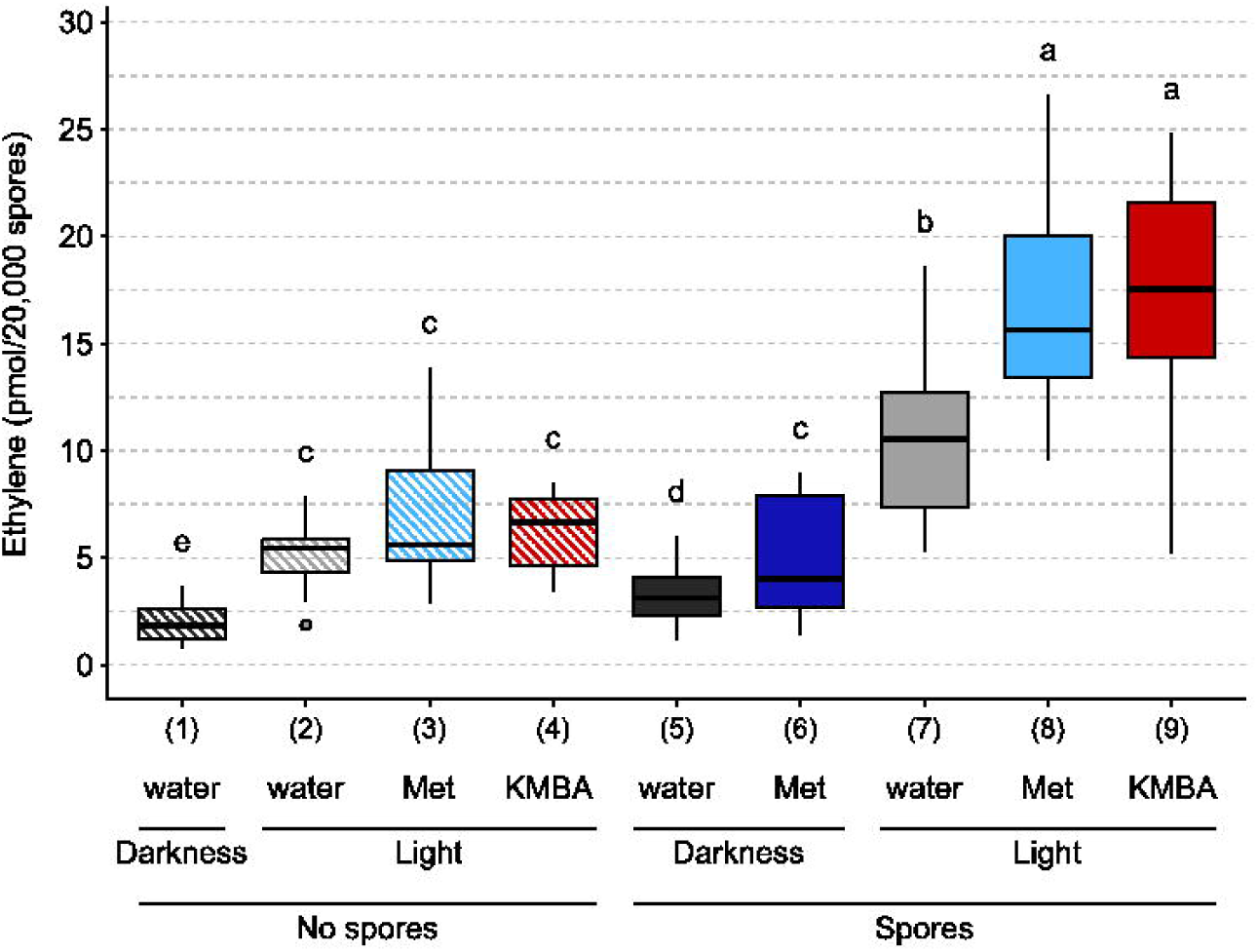
Ethylene production by *R. irregularis* in response to different treatments. 20,000 spores were germinated for three days in the dark, in the presence or absence of 10 mM methionine (Met) or 1 µM α-keto γ-methylthiobutyric acid (KMBA). Tubes were then sealed with a gas-tight stopper and exposed to light or darkness for 24h. One mL of the headspace gas was then analysed by gas chromatography. Different letters indicate different statistical groups (pairwise Kruskal-Wallis test with FDR correction, *P* < 0.05)

To investigate whether the KMBA pathway for ethylene synthesis was used by *R. irregularis*, we tested the effects of adding methionine into the incubation medium. In the dark, methionine addition increased ethylene production by 55 % (Fig 3, conditions 5 and 6). While in the light, methionine addition did not increase ethylene production in the absence of spores (Fig 3, conditions 2 and 3), this addition in the presence of spores increased ethylene production by 56 % (Fig 3, conditions 7 and 8). We then tested the effects of added KMBA. In the light and in the absence of spores, KMBA did not yield ethylene (Fig 3, conditions 2 and 4). In contrast, in the presence of spores, KMBA addition stimulated an ethylene production (Fig 3, conditions 7 and 9) similar to the methionine treatment.

Finally, we analysed by LC/MS the presence of KMBA in the exudates of spores germinated in the presence or not of methionine. This experiment was carried out in the dark to avoid light oxidation of KMBA. We could not detect the presence of KMBA in samples without methionine whereas we detected a strong KMBA signal in the methionine-treated samples, at a retention time of 3.7 min in MRM mode (*m/z* transitions 147 > 46 and 147 > 99) (S3 Fig).

Genomic sequences of Glomeromycotina were analysed for the presence of genes associated with the different ethylene biosynthesis pathways. The KMBA pathway requires the deamination of methionine into KMBA, which can be mediated by any transaminase, and the oxidation of KMBA into ethylene can be carried out non-specifically by peroxidases, or chemically [49,62,63]. Therefore, sequence analyses are not suitable to investigate the existence of this pathway. To look for genes associated with the major ACC pathway used by plants, AtACS1, AtACS8 and AtACS7 were chosen as representative of the three ACS clades described in Arabidopsis [64]. In the three cases, a tyrosine aminotransferase (GLOIN_2v1675208) and a pyridoxal phosphate-dependent transferase (GLOIN_2v1486204) were found as the best hits in *R. irregularis* genome, with only a maximum of 24% identity at the amino acid level with the query sequence. We also used AtACO2, AtACO1 and AtACO5 as representative of the three ACC oxidases (ACO) clades [65]. With these three queries, only hypothetical proteins with less than 26% of identity were found in *R. irregularis* datasets. To look for genes of the EFE pathway, BLAST analysis using the *Penicillium digitatum* EFE protein as query allowed the identification of two isoforms of a protein annotated as Fe (2+) and 2-oxoglutarate (2OG)-dependent dioxygenases in *R. irregularis* (GBC41587.1 and GBC41586.1). However, the similarity with the *P. digitatum* protein was very low (23-24% of identity).

## Discussion

Germinated spore exudates of AM fungi are known to trigger a number of symbiotically relevant responses in host root cells [66–68], indicating that isolated AM fungal spores release molecular signals within a few days of incubation. GSE form a matrix of relatively low complexity, which favours sensitive compound detection through mass spectrometry. We thus started our study by analysing GSE for the presence of a wide variety of phytohormones. The presence of iPR and IAA was unambiguously demonstrated by a combination of MRM and HRMS analyses, in GSE samples obtained from relatively small amounts of fungal material (10,000 spores). Although the presence of cytokinin-like compounds and of IAA was already suspected in AM fungi [54,55], the present study is to our knowledge the first conclusive report for the release of these two phytohormones by an AM fungus. It is of course possible that other phytohormones are present in low amounts in GSE, and have escaped detection despite the high sensitivity of the MRM approach (see S1 Table for detection limits).

Analysis of spore extracts was undertaken as a complementary approach to look for hormones that would not be released into GSE. This allowed the detection of MRM signals corresponding to GA_4_, at a retention time very close to that of the standard. Despite pre-purification, the fractions of spore extracts remain a complex matrix, which can induce slight shifts in chromatographic retention times. Using a GA_4_ standard to spike our sample allowed us to observe that the retention times recorded in the spore sample for both MRM transitions were identical to those of the standard in this matrix (Fig 2). Unfortunately, complex matrices decrease the sensitivity of HRMS analysis. We were thus unable to confirm the identity of this compound through accurate *m/z* determination. Our preliminary observations are however consistent with the previous bioassay-based detection of gibberellin-like compounds in AM fungi [54]. Given that gibberellins can be synthesised (and were actually discovered) in other fungi, it would make evolutionary sense to find them also in AM fungi

In contrast with Esch *et al.* [53], we did not detect the presence of ABA or glycosylated ABA. This difference might be due to the use of different fungal materials. The study of Esch *et al*. was carried out on an unspecified species of the genus *Rhizophagus* (formerly *Glomus*), and perhaps more importantly, the analysed material consisted of extraradical mycelium and spores obtained non axenically after several weeks of culture in pots. Furthermore, in this study, ABA detection was based on indirect ELISA tests, which likely differ from mass spectrometry in terms of sensitivity and specificity.

The detection of an auxin, a cytokinin and a gibberellin in *R. irregularis* does not mean that this fungus is actually able to synthesize these molecules. They could have been produced by the host plant (here the hairy roots of *Daucus carota*), imported by the fungus during the symbiotic exchanges between the two partners and stored in spores. Isotopic labelling experiments could be used to investigate the biosynthetic origin of these hormones, but the difficult incorporation of labelled precursors into fungal cells would likely limit the effectiveness of this approach.

Unlike the above hormones, there is no doubt about the fungal biosynthetic origin of ethylene. *De novo* ethylene production was measured over a period of 24 h and could be stimulated by the addition of methionine, a metabolic precursor. The addition of methionine strongly enhanced the synthesis of KMBA by the fungus (Fig S3) and acted synergistically with exposure to light to promote ethylene production (Fig 2). Light- and methionine-dependency are characteristic features of the KMBA pathway described in other fungi [47]. Hence, although we cannot rule out the existence of additional ethylene biosynthetic pathways, our biochemical data support the KMBA pathway as being involved in the synthesis of ethylene in *R. irregularis*. Interestingly, KMBA-derived ethylene was also demonstrated in the ectomycorrhizal fungi *Tuber brochii* and *T. melanosporum* [48]. To further investigate the existence of alternative pathways for ethylene synthesis in *R. irregularis*, we analysed Glomeromycotina genomic sequences. BLAST analyses failed to identify genes with a high homology with those involved in the classical ACC pathway found in plants. Similarly, the best hits obtained when searching for a fungal EFE exhibited limited identity with the query sequence. The fact that enzymes in the EFE family can be involved in the biosynthesis of a multitude of products [69] sheds further doubt on the role of their *Rhizophagus* homologs in ethylene synthesis. Further studies would be necessary to formally exclude the existence of these two biosynthetic pathways in *R. irregularis*, but in view of these initial investigations, their existence seems unlikely.

The observation that iPR, IAA and ethylene are released by germinating spores into their environment is consistent with a signalling role in the AM symbiosis. A similar hypothesis was proposed by Le Marquer *et al*. [56] with CLE peptides, another type of plant hormone potentially produced and excreted by AM fungi. First, if these hormones are still released at late stages, they could contribute directly to changes in hormonal contents in mycorrhizal plants. For example, AM colonization has been shown to increase auxin concentration in roots of *M. truncatula, Zea mays* and *Glycine max* [55,70,71]. In tomato roots, the expression of the auxin-dependent reporter DR5-GUS was higher in arbuscule-containing cells than in the surrounding cells [26]. This higher auxin concentration could be partly due to the AM fungus exudation and exportation to root tissues. Second, the release of these hormones by AM fungi could have profound effects on the symbiosis itself, such as the positive effects observed upon auxin treatment [72]. These hormones could also act through a modulation of plant development. For example, the simultaneous exudation of IAA and Myc-LCOs by the fungus could have synergistic effect on lateral root formation, as shown by Buendia *et al.* [73] with exogenous treatments on *Brachypodium distachyon*. The possible effects of ethylene released by the fungus are more difficult to predict. In arbuscular mycorrhiza, ethylene is mostly described as a negative regulator [30,66,74,75]. This conclusion was mainly drawn from studies using plant mutants disturbed in the production or perception of ethylene. As already proposed, the negative downstream effect of these mutations on AM symbiosis may also result from crosstalks with additional phytohormones and not directly from modifications in ethylene signalling *per se* [35]. Whether through direct or indirect mechanisms, the production of ethylene by AM fungi could serve to prevent excessive colonization of the root system. It is also important to note that ethylene inhibition of AM symbiosis was shown to be concentration-dependent [35] and that in some cases, a low ethylene concentration was able to stimulate root colonization [36].

In addition to hormonal signalling to the plant, it is also possible that AM fungi use phytohormones to regulate their own development. In support of this hypothesis, candidate genes putatively encoding ethylene and cytokinin receptors were recently identified in the genome of *R. irregularis* [76] and await functional characterization. This study was focused on histidine kinases, and thus does not exclude the existence of other types of receptors for other phytohormones in AM fungi. Indeed, a variety of hormones can affect AM fungal development *in vitro* [12–14,36,77,78]. It can also be noted that phytohormone exudation by plant roots is not restricted to strigolactones, and has also been reported for auxin, abscisic acid, jasmonate, salicylate and a cytokinin [79–82]. Bringing together these observations, it is tempting to speculate on the exchange of several hormonal signals in both directions during AM symbiosis, in addition to the well-known effects of phytohormones as internal regulators of plant physiology. Interestingly, the use of a common language has been reported in diverse contexts of host-microbe interactions. For example, plant bacterial pathogens produce cytokinin and have evolved a corresponding receptor [83], and gut bacteria produce and possess sensors for neuroendocrine hormones that were once thought to be specific of their host [84,85]. Plants have lived with AM symbionts since they colonized land, and the molecular language underlying this long-standing and intimate relationship is only beginning to be unravelled. Deciphering the hormonal biosynthesis and perception pathways in AM fungi will certainly help to understand how this common language developed through evolution.

## Acknowledgments

We thank Cyril Libourel, Marielle Aguilar and Helene San Clemente for their help with statistical analyses. We thank Maryne Laigle, Virginie Durand and Thibaut Perez for their help with mass spectrometry analysis.

Support for mass spectrometry analyses was provided by the ICT-Mass spectrometry and MetaboHUB-MetaToul-AgromiX Facilities.

## Supporting information

**S1 Table.** List of molecules analyzed by LC-MS in highly sensitive MRM mode The hormone family, molecule name, abbreviation, formula, retention time, preferential detection mode, precursor and product ions for MRM analysis and limit of detection are indicated. Grey lines correspond to theoretical fragmentation values when standards were not available [86,87].

**S1 Fig.** High resolution mass spectra (HRMS) of iPR in positive mode (A) and (B) HRMS spectra of iPR standard (300 nM), at 5.78 min. (C) and (D) HRMS spectra of 250,000 *R. irregularis* GSE at 5.78 min. (A) and (C), MS spectra. (B) and (D) MS/MS (*m/z* 336.1666 +/− 0.5 Da) spectra.

**S2 Fig**. High resolution mass spectra (HRMS) of IAA in positive mode (A) and (B) HRMS spectra of IAA standard (300 nM) at 6.19 min. (C) and (D) HRMS spectra of 250,000 *R. irregularis* GSE, at 6.19 min. (A) and (C) MS spectra. (B) and (D) MS/MS (*m/z* 176.0705 +/− 0.5Da) spectra.

**S3 Fig.** KMBA detection in multiple reaction monitoring (MRM) mode (A) Structure and fragmentation pattern of KBMA. (B) UPLC-MRM-MS chromatogram of KMBA in negative mode. Top (blue lines): External standard (500 nM) of KMBA. Middle (red lines): GSE produced by 20,000 spores treated with 10 mM methionine. Bottom (green lines): GSE produced by 20,000 untreated spores. Plain lines are for *m/z* transition 147 > 47. Dashed lines are for *m/z* transition 147 > 99. Signal intensity is displayed in counts per second (cps).

## Notes

### Competing Interest Statement

The authors have declared no competing interest.

### Summary of Updates

Minor lexical and bibliographical revisions.

