## Supplementary figures and images for "Phytohormone production by the arbuscular mycorrhizal fungus *Rhizophagus irregularis*"

### Supplementary figure 1

MS and MS/MS spectra of iPR

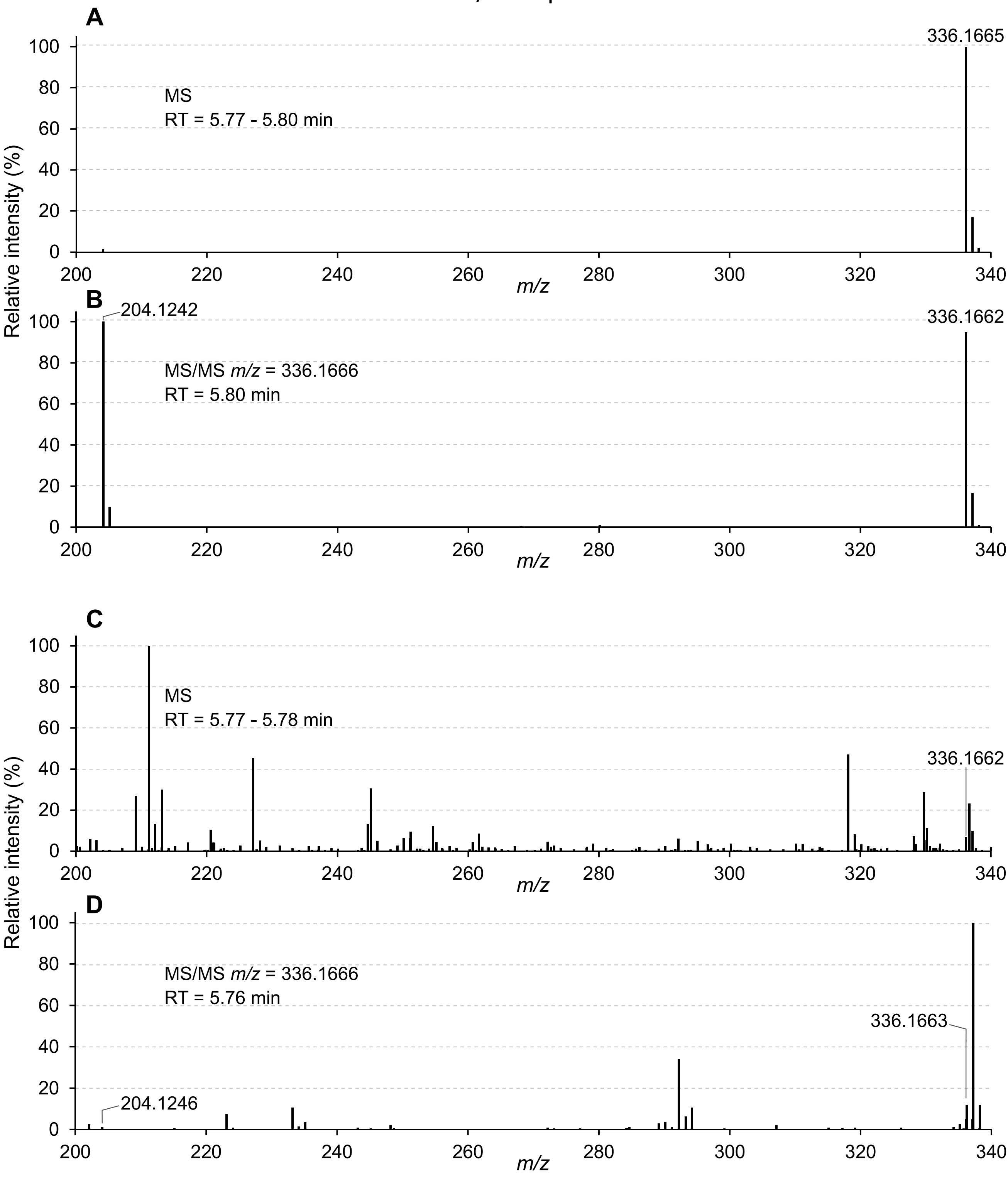

### Supplementary figure 2

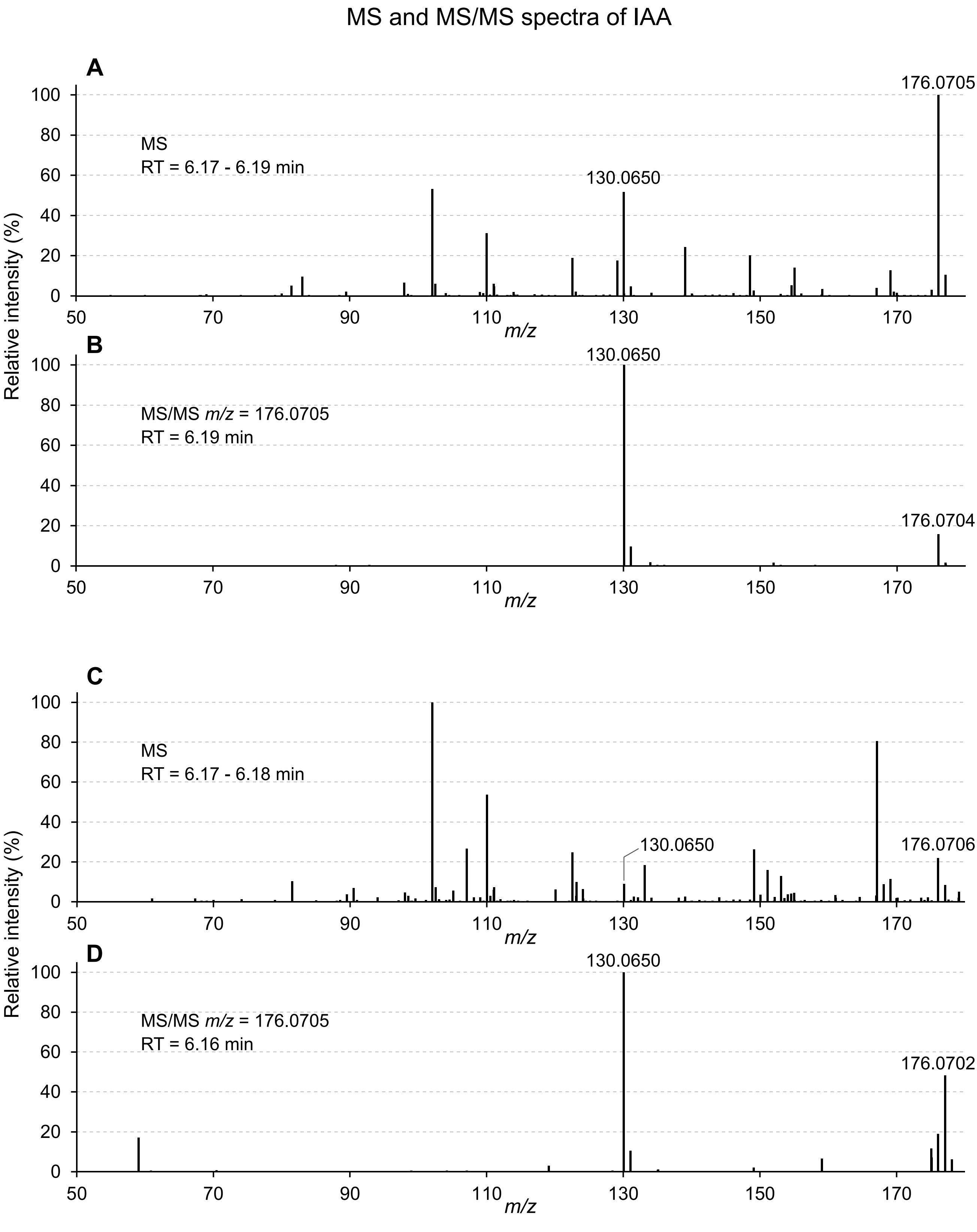

### Supplementary figure 3

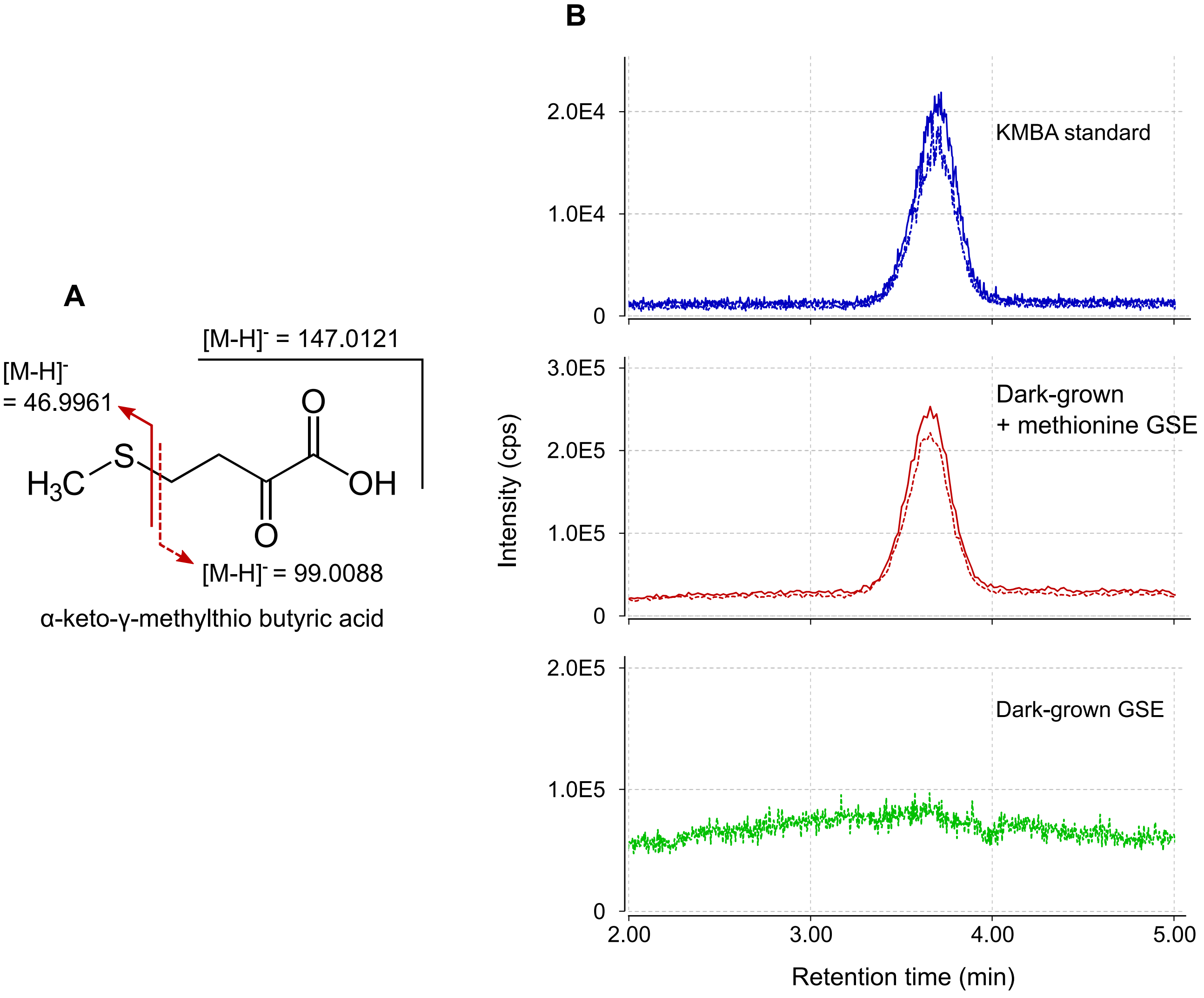
